# Magnetite biomineralization in Magnetospirillum magneticum is regulated by a switch-like behavior in the HtrA protease MamE

**DOI:** 10.1101/047555

**Authors:** David M. Hershey, Patrick J. Browne, Anthony T. Iavarone, Joan Teyra, Eun H. Lee, Sachdev S. Sidhu, Arash Komeili

## Abstract

Magnetotactic bacteria are aquatic organisms that produce subcellular magnetic particles in order to orient in the earth’s geomagnetic field. MamE, a predicted HtrA protease required to produce magnetite crystals in the magnetotactic bacterium *Magnetospirillum magneticum* AMB-1, was recently shown to promote the proteolytic processing of itself and two other biomineralization factors *in vivo*. Here, we have analyzed the *in vivo* processing patterns of three proteolytic targets and used this information to reconstitute proteolysis with a purified form of MamE *in vitro*. MamE cleaves a custom peptide substrate with positive cooperativity, and its auto-proteolysis can be stimulated with exogenous substrates or peptides that bind to either of its PDZ domains. A misregulated form of the protease that circumvents specific genetic requirements for proteolysis causes biomineralization defects, showing that proper regulation of its activity is required during magnetite biosynthesis *in vivo*. Our results represent the first reconstitution of MamE’s proteolytic activity and show that its behavior is consistent with the previously proposed checkpoint model for biomineralization.

## Introduction

Magnetotactic bacteria assemble iron-based magnetic crystals called magnetosomes into chains within their cells allowing them to passively align in and navigate along magnetic fields(c1, c2). Understanding the mechanism of biomineralization in these organisms can provide novel strategies for manipulating transition metal-based nanomaterials *in vitro(c3–c5)*. Genetic analyses have identified a set of genes, called biomineralization factors, whose deletions disrupt or eliminate magnetite crystal formation(c6–c10). Two of these, *mamE* and *mamO*, encode predicted trypsin-like proteases required to produce magnetite in the model magnetotactic organism, *Magnetospirillum magneticum* AMB-1(c6). Disrupting either gene abolishes the formation of magnetite crystals without disturbing the production of their surrounding membrane compartment, showing that each protein is required for initiating magnetite biosynthesis(c6, c11, c12).

Cells with a catalytically inactive (*E^PD^*) allele of *mamE* show an intermediate biomineralization phenotype in which they produce small magnetite particles. Wild-type AMB-1 has a distribution of crystal sizes centered at 50-60nm in diameter, but the size distribution in the *E^PD^* cells is centered at ~20nm. Interestingly, ~97% of the crystals in the *E^PD^* strain are smaller than 35nm, the point above which magnetite particles become paramagnetic and can hold a stable magnetic dipole. The correlation between mineral sizes in the *E^PD^* strain and the superparamagnetic to paramagnetic transition point lead to a model predicting that cells produce small superparamagnetic crystals until an unknown signal activates MamE, promoting maturation to paramagnetic particles(c13).

MamE is a member of the HtrA/DegP family of trypsin-like proteases, a ubiquitous family of enzymes that controls various aspects of protein quality control(c14). The family is characterized by domain structures consisting of an N-terminal trypsin-like domain and one or two C-terminal postsynaptic density 95/discs large/zonula occludens-1 (PDZ) domains(c15–c17). Structural and mechanistic investigations indicate that the PDZ domains regulate proteolysis by promoting assembly and activating the protease domain by binding to extended peptide motifs(c18–c23). Recently, MamE was found to promote the *in vivo* proteolytic processing of itself, MamO, and another biomineralization factor named MamP, in a manner that required the predicted MamE active site (Fig. 1A)(c24–c26). Although MamO was also required for these proteolytic events, this effect did not require the its predicted protease active site. Subsequent structural analysis showed that MamO’s protease domain was locked in an inactive state and incapable of catalysis, suggesting that it played a non-catalytic role in activating MamE(c24).

**FIGURE 1.**
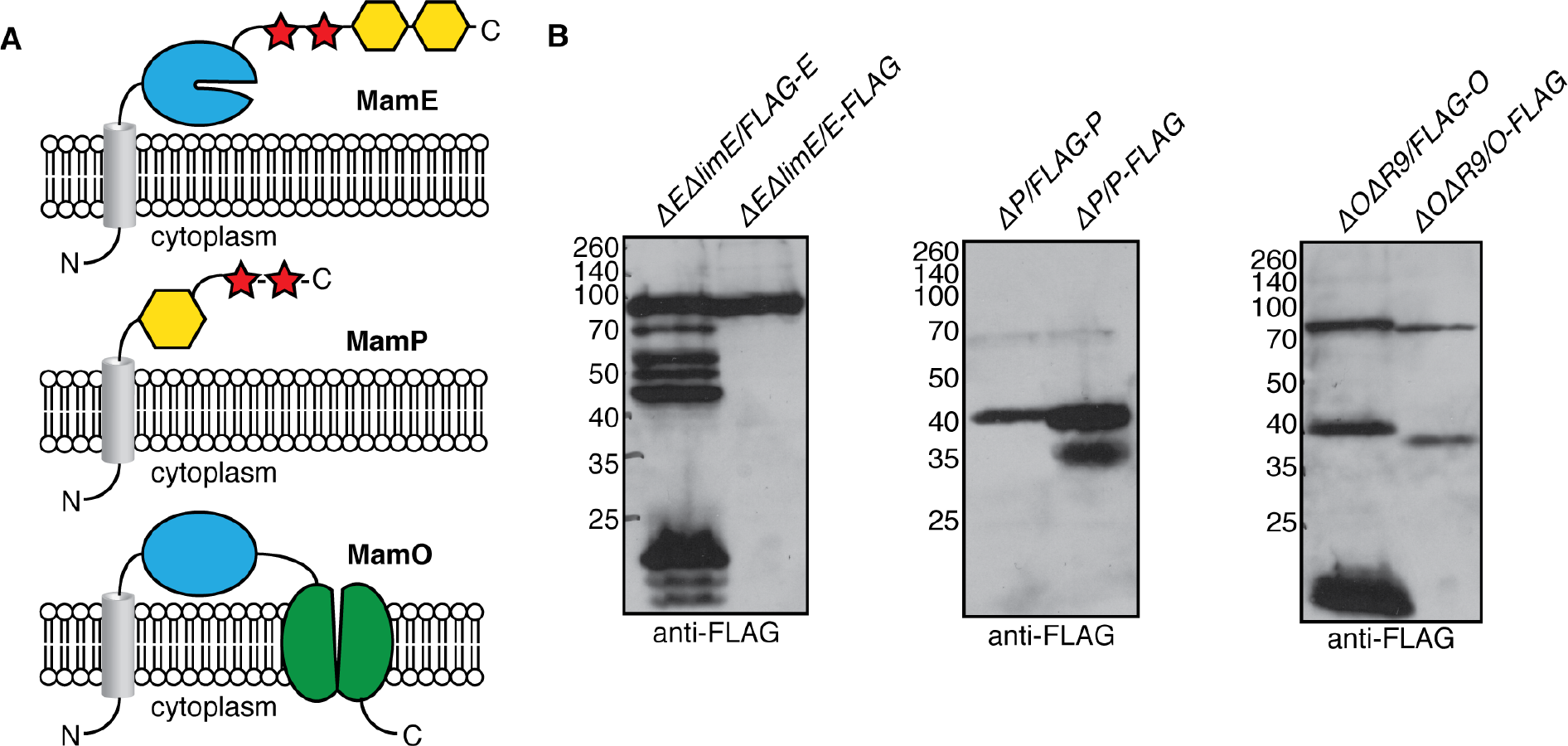
*In vivo* proteolytic processing of MamE, MamP and MamO. *A*, Predicted domain structures of the three proteolytic targets. Grey cylinder: transmembrane helix; blue: trypsin-like domain; red: *c*-type cytochrome; yellow: PDZ domain; green: TauE domain. The MamO protease domain is represented differently due to its assignment as an inactive pseudo-protease. *B*, Proteolytic processing patterns observed through epitope tagging. The fragments were observed in each of at least four independent experiments with the exception of the 20kDa N-terminal fragment of MamE and the 17kDa N-terminal fragment of MamO, which varied dramatically between experiments.

Despite the genetic evidence for MamE’s role in proteolysis, its activity has not been confirmed directly using purified components. Studies aimed at understanding the catalytic activity of MamE and its regulation have been hindered by an inability to obtain recombinant protein. Here, we have characterized MamE-dependent proteolysis in detail and identified a number of regulatory mechanisms. Developing a method to purify MamE and analyzing *in vivo* proteolytic patterns of each target facilitated reconstitution of MamE-dependent proteolysis *in vitro*. Detailed analysis of its catalytic activity suggests a switch-like model in which the basal state of the protein is an inactive form that can be turned on through a number of routes. A constitutively active allele of MamE disrupts biomineralization, confirming that properly regulated proteolysis is critical to magnetosome formation.

## Results

### *Mapping MamE-dependent cleavage patterns* in Vivo

To determine the context of MamE-dependent proteolysis, we analyzed the cleavage patterns of MamE, MamP and MamO using an epitope tagging approach. N or C-terminally-tagged alleles of each gene were added to their respective deletion strains. Each of the tagged alleles complemented the biomineralization defects of the deletions with the exception of the C-terminally tagged *mamO* allele *(O-FLAG)*. For MamE, a number of N-terminal proteolytic fragments but no C-terminal fragments were observed (Fig. 1B), indicating that short segments are sequentially removed from the C-terminus. One N-terminal and two C-terminal bands of MamP were observed, indicating the removal of a fragment containing the membrane anchor from the predicted soluble region (Fig. 1B). For MamO, a full-length and a shorter band were observed for both the N and C-terminally tagged proteins, predicting an internal MamE-dependent cleavage site resulting in two stable fragments (Fig. 1B). The levels and presence of the small (i.e. ~20 kDa) bands appearing in blots of N-terminally tagged MamE and MamO were highly inconsistent. This could suggest that short segments are removed from the N-terminus of each protein to produce unstable fragments that are quickly degraded, but due to their inconsistency we have not focused on them in our analysis of the processing pattern.

### Identification of a putative cleavage motif in MamO

We reasoned that one strategy for reconstituting MamE’s proteolytic activity could be to design a substrate based on an *in vivo* cleavage motif in one its targets. MamO’s processing pattern indicated the presence of a ~37kDa N-terminally tagged fragment containing a MamE-dependent cleavage site at its mature C-terminus. Cell pellets from magnetic cultures of the *ΔOΔR9/FLAG-O* strain were used for biochemical fractionation (Fig. 2A). Enzymatic lysis and sonication followed by a low speed (8000 × g) centrifugation were used to isolate material associated with the dense magnetite particles. A number of detergents were tested for their ability to dissolve the MamO fragments. Most classes of detergents were ineffective or only partially effective in the initial solubilization step, but lipid-like zwitterionic detergents including lauryldimethylamine oxide (LDAO) and FosCholine-12 efficiently extracted the fragments from the membrane. Although these detergents disrupted binding to the α-FLAG affinity resin, once the initial extraction step was complete, the detergent requirements to maintain solubility became less stringent.

Based on the solubility information, the low speed pellet was pre-washed with CHAPS before extracting MamO fragments from the membranes with FosCholine-12. In order to facilitate binding to the affinity resin, FosCholine soluble material was loaded on an anion exchange column and exchanged to the detergent DDM by extensive washing before eluting with salt. This fraction was then used as the input for an α-FLAG affinity isolation to yield a final fraction enriched in N-terminal fragments of MamO (Figs. 2A and Fig. 2B). The concentrated α-FLAG elution was separated on an SDS-PAGE gel, stained with colloidal Coomassie Blue and the region around 37kDa was excised. After performing in-gel trypsin digestion, peptides were extracted and concentrated for LC-MS/MS analysis. A number of peptides from the MamO sequence were consistently detected, and, as expected, they mapped almost exclusively to the protease domain of MamO (Fig. 2C). In all of the samples, the protein sequence coverage dropped off sharply in the linker between the protease and TauE domains (Fig. 2C). A peptide with the sequence GSATAPGQPQTQTV was routinely detected at the C-terminal edge of the peptide coverage (Fig 2D). This peptide results from a predicted tryptic cleavage on the N-terminus but has a non-tryptic C-terminus, which suggests that it contains the C-terminal sequence of the mature MamO protease domain.

**FIGURE 2.**
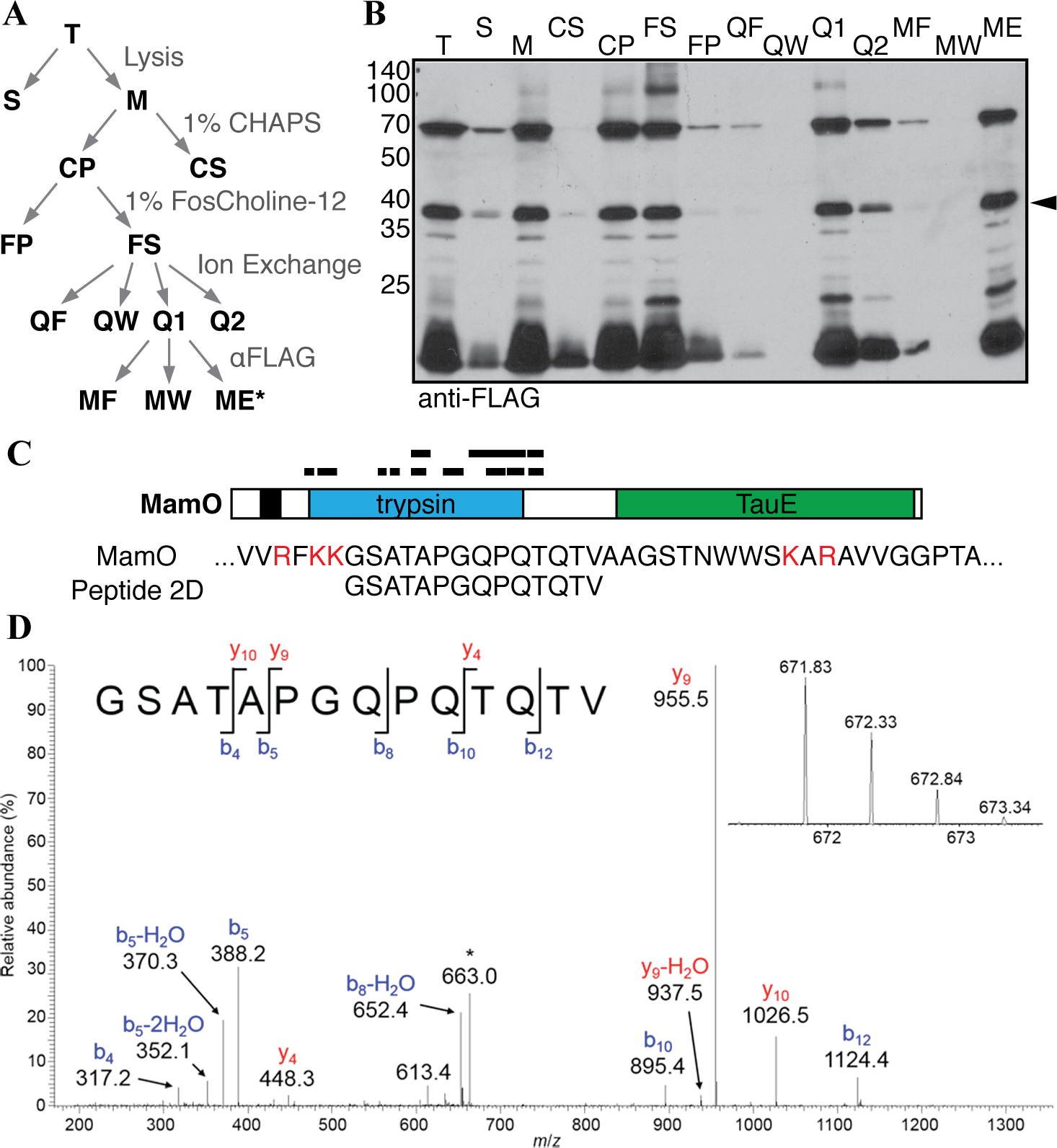
Biochemical fractionation to enrich N-terminal MamO fragments. *A*, Schematic of the enrichment procedure. *B*, Immunoblot of each fraction from A. The predicted protease domain fragment is marked with a carat. *C*, Peptides identified in a representative LC-MS/MS analysis. The red letters in the MamO sequence represent predicted tryptic cleavage sites. The coverage pattern is characteristic of analyses for three separate preparations. *D*, Tandem mass spectrum from collision-induced dissociation of the [M+2H]^2+^ ion of the peptide, GSATAPGQPQTQTV, corresponding to amino acid residues 273-286 of MamO. The inset shows detail for the isotopically resolved, unfragmented peptide precursor ion. The fragment ion at *m/z* = 663 (denoted by the asterisk) is due to precursor ion that has undergone neutral loss of a molecule of water. This peptide was detected in each of three biological replicate experiments.

### Direct proteolysis of the MamO cleavage motif

We expressed and purified a form of MamE with the N-terminal membrane anchor removed in order to study proteolysis of the *in vivo* cleavage motif identified in MamO. The recombinant protein was targeted to the *E. coli* periplasm to allow for heme-loading of the c-type cytochrome motifs (Fig. 3). We designed a fluorogenic peptide containing the 8 residues of the putative cleavage motif identified in MamO (Fig. 2) flanked by an upstream fluorophore and a downstream fluorescence quencher. Normally, the peptide has low fluorescence due to interaction between the fluorophore and quencher. If the peptide is cleaved, the two fragments will separate, resulting in an increase in fluorescence. Upon addition of the O1 peptide to purified MamE, there was a linear increase in fluorescence. Importantly, the MamE^S297A^ protein did not alter the fluorescence indicating the signal was due to serine protease activity from MamE (Fig. 4A). MamE hydrolyzed the O1 peptide with a *k_cat_* of 0.64 ± 0.03 min^−1^ and a *K_M_* of 6.1± 0.5 μM. Interestingly, the reaction was positively cooperative, displaying a Hill coefficient of 1.5 ± 0.1 (Fig. 4B) (c18, c35). These values are similar to those reported for cleavage of peptide substrate by other trypsin-like proteases and confirm that MamE can efficiently cleave the motif identified in MamO(c36). Combined with the *in vivo* analysis these results confirm that MamO is a direct proteolytic target of MamE.

**FIGURE 3.**
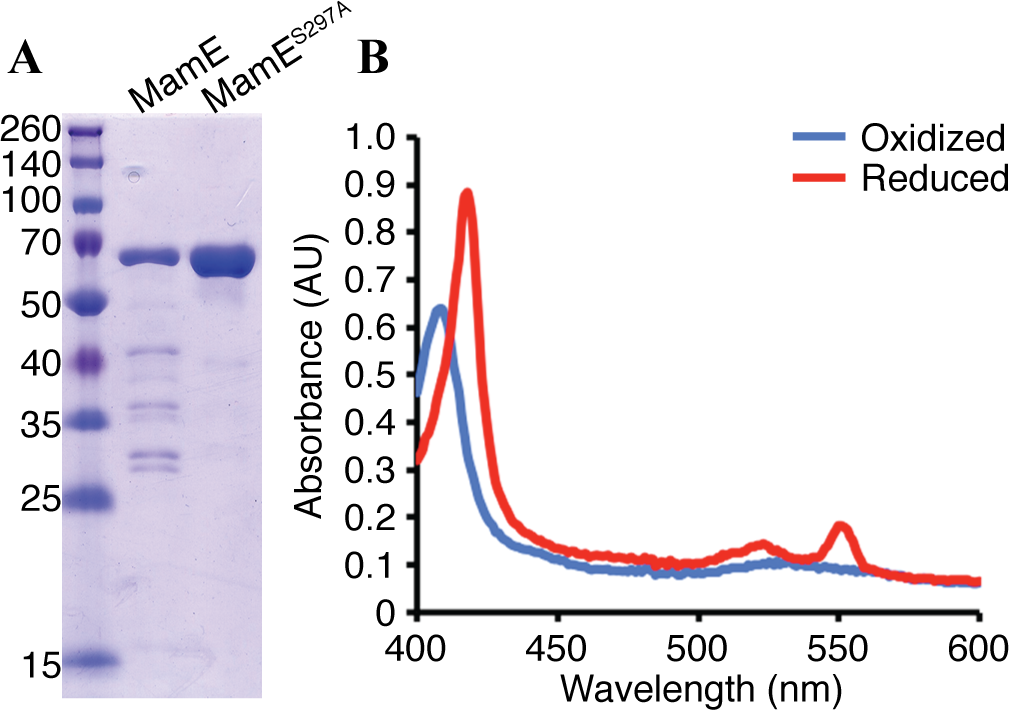
Purification of MamE. *A*, MamE and MamE^S297A^ (residues 108-728) were purified as a fusion to an N-terminal 6xHis and C-terminal strep tag. *B*, Absorbance spectrum of MamE^S297A^ in the oxidized (blue) and dithionite reduced (red) forms.

**FIGURE 4.**
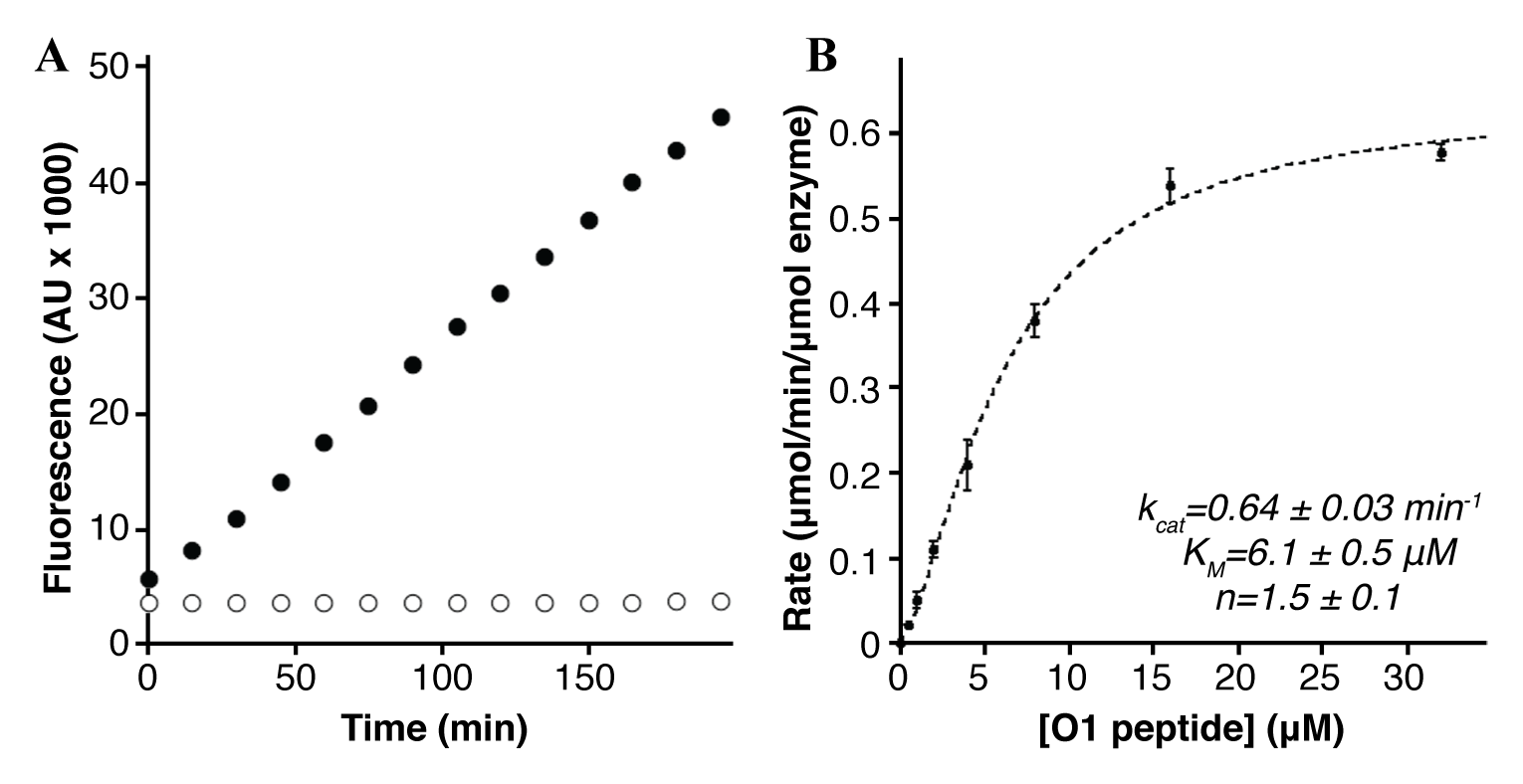
Cleavage of the MamO1 fluorogenic substrate. *A*, Linear increase in fluorescence upon addition of 20μM MamO1 peptide to 200nM MamE (black circles). No increase is seen with peptide addition to MamE^S297A^(white circles). *B*, Steady-state kinetics of O1 cleavage by MamE. The dotted line represents a fit to the Hill form of the Michaelis-Menten equation. Error bars represent the standard deviation from three technical replicates. The plot is characteristic of the data seen in five biological replicates.

### Reconstitution of MamE auto-proteolysis

Analysis of MamE processing in AMB-1 along with the extensive auto-proteolysis during its expression in *E. coli* suggested that MamE is capable of auto-proteolysis. However, purified MamE was relatively stable such that after an hour of incubation at 30°C, nearly all of the protein remained intact (Fig. 5A). The positive cooperativity observed for the steady-state kinetics of O1 peptide cleavage indicated that MamE’s catalytic activity could be stimulated by substrates. We reasoned that this mode of regulation might also lead to peptide-induced activation of autocleavage. Indeed, the MamO1 peptide stimulated degradation of full-length MamE in a dose-dependent manner, confirming that MamE’s activity can be stimulated by the presence of substrate (Fig. 5A).

**FIGURE 5.**
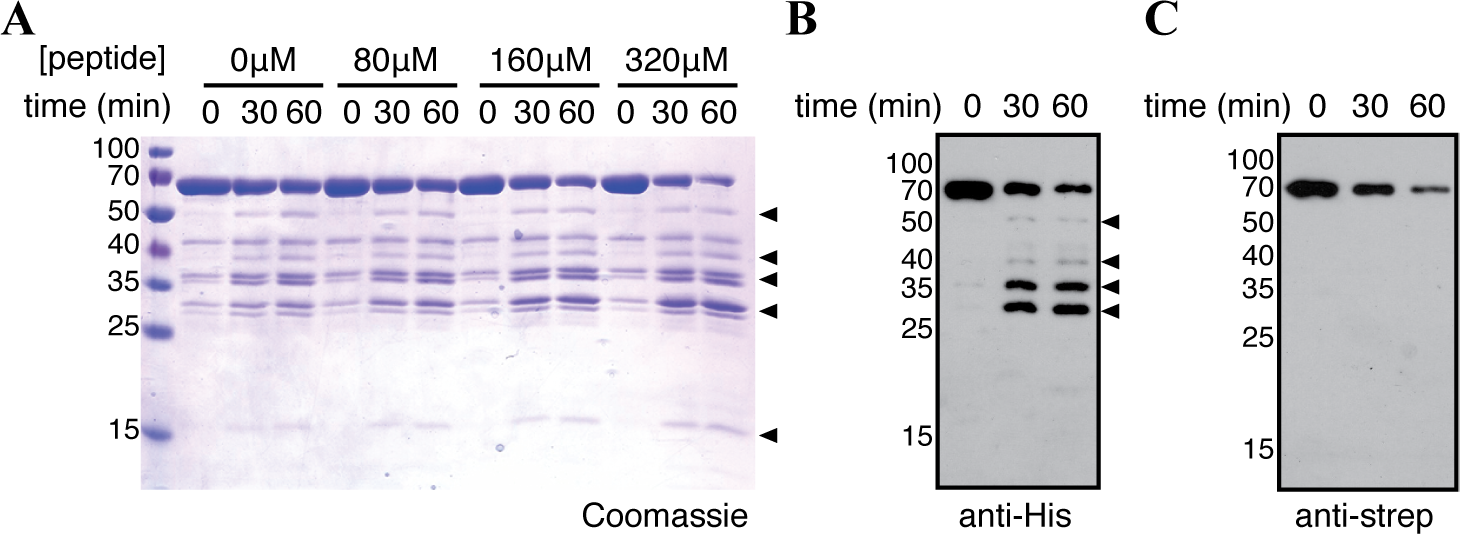
Reconstitution of MamE auto-processing. *A*, The MamO1 substrate induces auto-cleavage of MamE. *B, C*, Processing patterns assessed by immunoblotting of a reaction containing 320μM O1 for the indicated tag on MamE. The experiment is representative of three biological replicates. Throughout, proteolytic fragments increasing during the assay are marked with carats.

Taking advantage of the distinct tags used to purify MamE, we examined the auto-cleavage fragmentation pattern by immunoblotting. Numerous truncated proteins were detected by blotting for the N-terminal 6xHis tag, the smallest of which is a ~27 kDa fragment presumed to be the protease domain (Fig. 5B). In contrast, only the full-length protein was seen when blotting for the C-terminal strep tag (Fig. 5C). The pattern indicates that the reaction proceeds via sequential removal of small fragments from the C-terminus. Furthermore, it matches the pattern seen by examining epitope-tagged alleles of MamE expressed *in vivo* and confirms the successful reconstitution MamE-dependent proteolysis *in vitro*.

### Activation through the PDZ domains

PDZ domains often regulate proteolytic activity by binding to extended peptide motifs(c19). Phage display has been a productive approach for identifying peptide ligands that bind to PDZ domains(c37–c40). Each of the MamE PDZ domains was purified and used as bait in phage display selections. Both bait proteins showed phage enrichment for specific particles with a library displaying peptides on the N-terminus of the coat protein, but no enrichment was observed in libraries displaying C-terminal fusions. This suggests that, unlike those associated with other HtrA proteases, the MamE PDZ domains do not display a preference for C-terminal peptides(c30, c41). Interestingly, phage selections for both domains showed a strong preference for internal regions (Table 3 and Table 4). However, a single clone dominated both pools making the identification of consensus motifs hard to interpret.

**Table 3.**
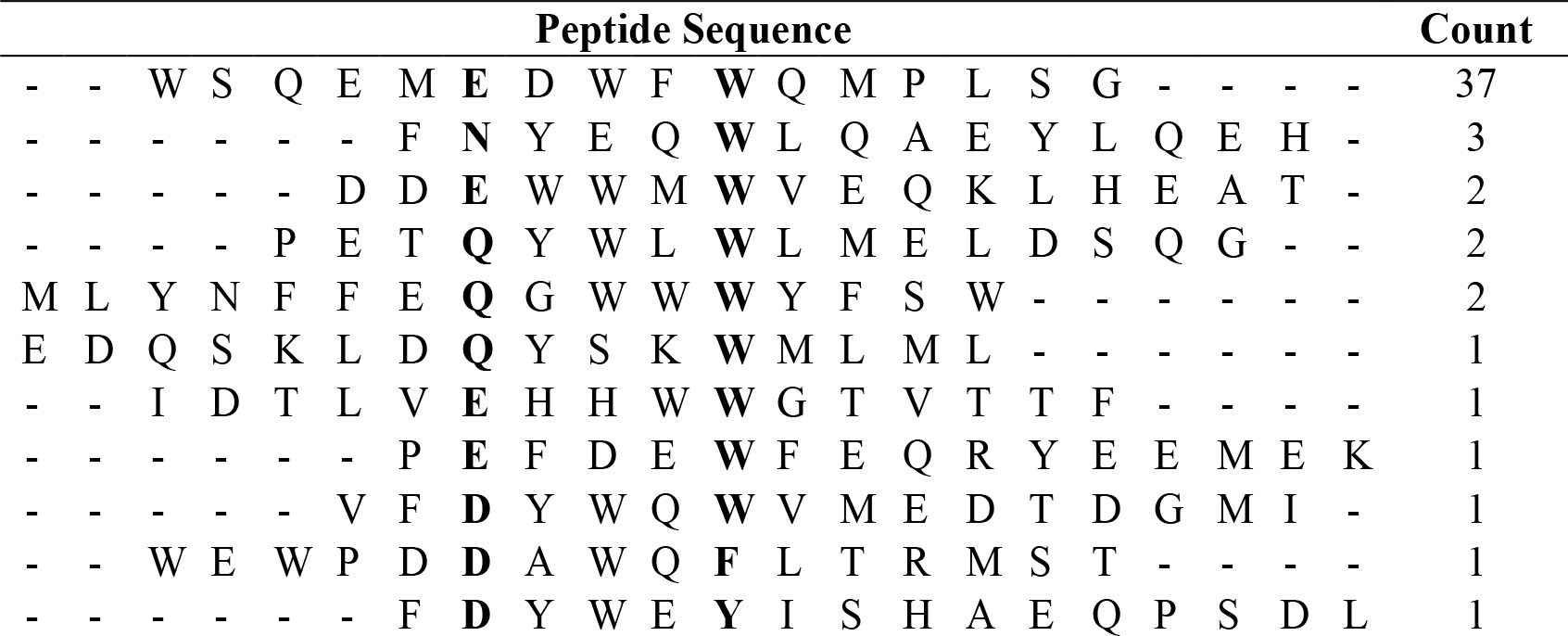
Peptide sequences from the phage display selections using MamE PDZ1. Each sequence was identified by isolating plaques from the enriched phage library and sequencing the gene for the major coat protein. The count represents the number of individual clones displaying the indicated peptide. The sequences are aligned based on the motif ΨxxxΩ, where Ψ is a hydrophilic residue and Ω is an aromatic residue. Bold type is used to mark the hydrophilic and aromatic residues around which the sequences were aligned.

**Table 4.**
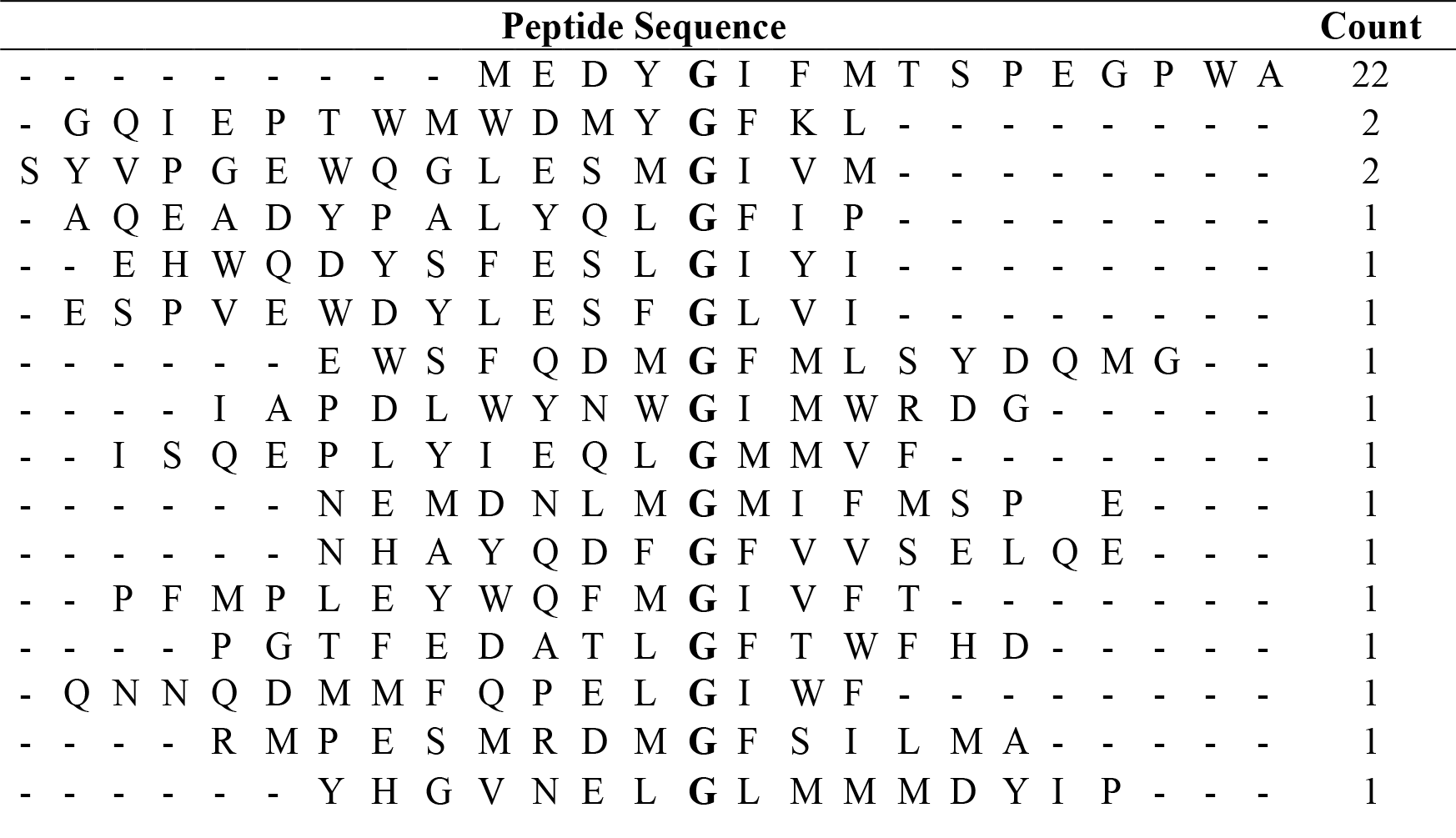
Peptide sequences from the phage display selections using MamE PDZ2. Each sequence was identified by isolating plaques from the enriched phage library and sequencing the gene for the major coat protein. The count represents the number of individual clones displaying the indicated peptide. The sequences are aligned based on the motif Ψ**ΦGΦΦΦ**, where Ψ is a hydrophobic residue and **Φ** is a hydrophilic residue. Bold type is used to mark the glycine residue around which the sequences were aligned.

Peptides corresponding to the sequence that dominated each selection were synthesized and labeled with a fluorescent dye. In addition, the C-terminal region of MamE containing only the PDZ domains (EP12) was purified and used to test for direct binding to the phage-derived ligands. Fluorescence anisotropy experiments demonstrated that both phage-derived peptides bind to the C-terminus of MamE (Fig. 6A). The PDZ1 peptide showed ~10-fold tighter binding than the PDZ2 peptide, but both affinities were comparable to those seen for other PDZ domains(c40). Addition of either peptide to full-length MamE, resulted in a dose-dependent activation of auto-processing (Fig. 6B). Importantly, the activation threshold for the PDZ2 peptide was higher than the PDZ1 peptide, mirroring the equilibrium binding data. Thus, ligand binding to either of MamE’s PDZ domains activates its auto-cleavage activity.

**FIGURE 6.**
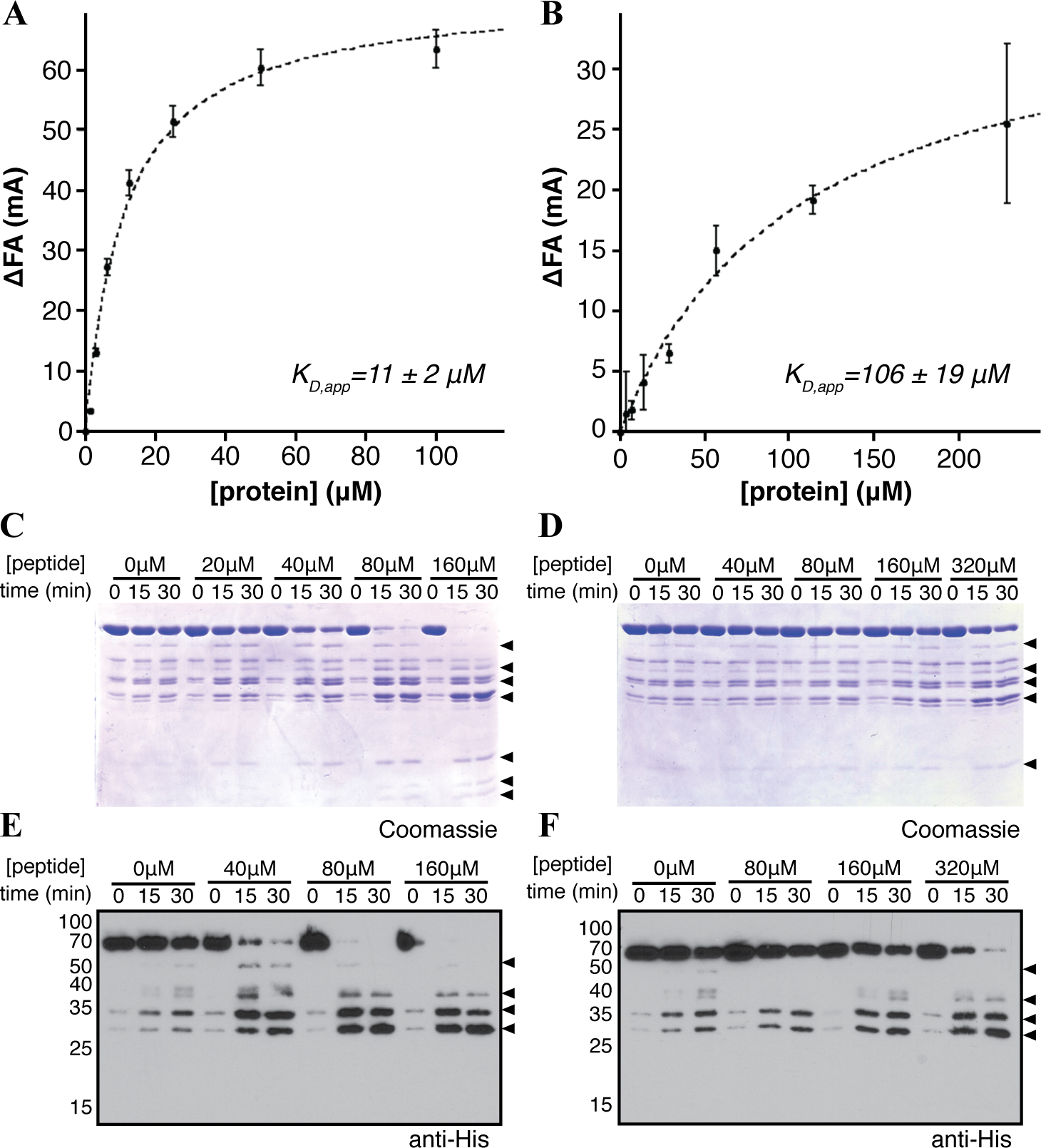
Peptide binding to the PDZ domains activates MamE. *A*, Fluorescence anisotropy showing binding of the EPDZ1^*^ peptide to the EP12 protein. The error bars represent the standard deviation from three technical replicates. The dotted line represents a fit of the data to a single-site binding model. *B*, Binding of the EPDZ2^*^ peptide to the EP12 protein. *C*, Activation of MamE auto-cleavage by the EPDZ2^*^ peptide. *D*, Activation of MamE auto-cleavage by the EPDZ2^*^ peptide. Each experiment was repeated a minimum of three times. *E*, Anti-His immunoblot of the reaction in *C. F*, Anti-His immunoblot of the reaction in *D*. Throughout, proteolytic fragments increasing during the assay are marked with carats.

### *Misregulation of MamE disrupts biomineralization* in Vivo

The switch-like activation of MamE’s catalytic rate suggested that its activity is carefully modulated during biomineralization. Mutating residue 192 (chymotrypsin numbering) in the oxyanion hole to proline increases basal cleavage rates in the model HtrA protease DegS(c18, c42). We introduced an allele with the analogous mutation of MamE (Q294P) into the *mamE* null strain (Fig. 7A). This strain displayed significantly less MamE and MamP, consistent with the expectation that the MamE^Q294P^ variant is a more active protease. Additionally, the *mamE^Q294P^* allele partially circumvents the requirement for *mamO* in promoting MamE-dependent proteolysis. MamE and MamP were not processed when the *mamE^WT^* allele was introduced into a strain lacking *mamE* and *mamO*. However, when the *mamEF^294P^* allele was introduced in the same background, proteolysis of MamE and MamP was restored, though processing of MamE does not reach to wild-type levels (Fig. 7B and Fig. 7C). The intermediate levels of MamE in the absence of MamO indicate that the *MamE^Q294P^* protein is not inherently unstable. These results show that *mamEF^294P^* produces a misregulated protease, leading to increased processing of biomineralization factors *in vivo*.

**FIGURE 7.**
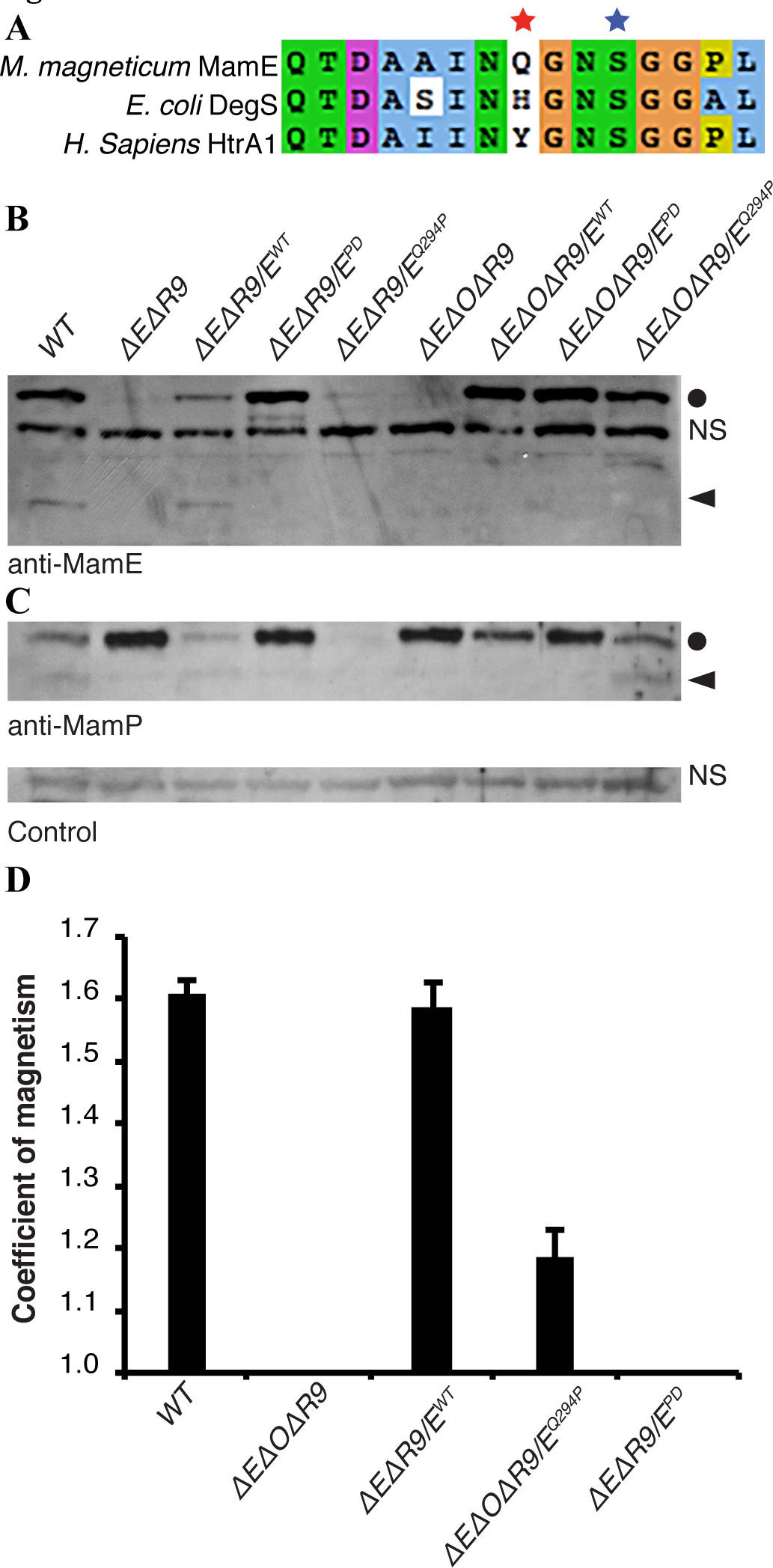
A constitutively active form of MamE disrupts biomineralization. *A*, Alignment of HtrA proteases. The red star marks position 192 in the oxyanion hole (residue 198 in DegS), and the blue star marks the catalytic serine nucleophile. *B* and *C*, Immunoblot analysis of AMB-1 lysates probed for MamE (B) and MamP (C). Circles mark full-length proteins and carats mark proteolytic fragments. NS marks nonspecific bands reacting with each antibody preparation. *D*, Magnetic response of AMB-1 cultures with the indicated genetic background. Biological replicates represent independent cultures of each strain and each measurement represents the average and standard deviation from three independent experiments.

We next examined the *mamE^Q294P^* allele for its ability to complement the biomineralization defects seen in the *mamE* null strain. While the *mamE^WT^* allele completely restores the magnetic response, the *mamE^Q294P^* strain has a significantly lower response (Fig. 7D). Thus, both the inactive *(mamE^PD^)* and misregulated forms *(mamEF^294P^)* of the protease disrupt biomineralization *in vivo*. The magnetic response of the *mamE^Q294P^* strain is higher than the negligible signal measured in the *mamE^PD^* strain, indicating that the biomineralization process is stalled at a different stage when MamE’s activity is unregulated. These results demonstrate that complete biomineralization of magnetosome crystals relies not only on the occurrence of MamE-dependent proteolysis but also careful regulation of the activity.

## Discussion

Biochemical principles underlying the biomineralization of magnetite by magnetotactic bacteria represent a model for understanding how biological molecules manipulate inorganic compounds(c43). Recent advances describing the genetic basis for this process have paved the way for mechanistic studies of the factors that promote mineral formation(c6). Through these efforts, the HtrA protease MamE has emerged as a central biomineralization factor in the model organism *Magnetospirillum magneticum* AMB-1. In addition to promoting crystal nucleation within the magnetosome lumen, MamE regulates a transition from small superparamagnetic crystals to full-sized paramagnetic particles. This crystal maturation phenotype was specifically linked to MamE’s putative protease activity suggesting a model where its catalytic activity controls crystal maturation(c13).

Here, we have studied MamE’s serine protease activity in detail. We have mapped the proteolytic patterns of three *in vivo* targets both at the domain level and, in one case, at the individual residue level. Using this information, we have reconstituted a number of aspects of MamE-dependent proteolysis with purified components. We show that MamE directly cleaves a motif from the linker between MamO’s protease and TauE domains in a positively cooperative fashion. Furthermore, we show that purified MamE has low basal activity, but that it can be activated in a number of ways including the presence of substrate and peptide binding to either of its PDZ domains. This behavior is consistent with a switch-like mode of regulation in which the protease requires activation by environmental cues. These results also show for the first time that MamE is a serine protease that is capable of directly degrading itself and other biomineralization factors.

The peptide-based substrate and ligands developed here shift MamE into the activated state *in vitro*, but the specific signals that activate the protein *in vivo* are not known. We previously showed that proteolysis of MamE, MamO and MamP requires the presence of MamO’s C-terminal ion transporter domain, suggesting that manipulation of the solute environment in the magnetosome regulates MamE’s activity(c24). However, the transport cargo(es) of MamO have not been identified, making the prediction of potential activating solutes difficult. In addition to its protease and PDZ domains, MamE contains a pair of *c*-type cytochrome motifs called magnetochrome domains that are commonly seen in magnetosome biomineralization proteins (Fig. 1A). MamE’s tandem magnetochrome region was previously purified and displayed a single midpoint redox potential of −32mV, consistent with the proposed roles of similar biomineralization factors in controlling the redox status of iron(c25, c26, c44). Though redox mediated activation presents an attractive model for MamE regulation, we have not observed any redox dependence for the rate of auto-proteolysis or steady-state cleavage of the MamO1 peptide.

Despite the uncertainty about the specific events that trigger its activation, the switch-like modulation of MamE’s activity *in vitro* suggested that allosteric regulation of its protease activity was required for proper crystal maturation. We developed an allele of MamE with a mutation reported to stabilize active forms of other HtrA proteases and showed that this allele no longer required *mamO* to promote proteolysis *in vivo*. Like the catalytically inactive form, this misregulated form of MamE had defects in crystal maturation(c18). These results confirm that both the active and inactive states are important during the process of magnetosome formation. Similar experiments with DegP in *E. coli* indicated that the proper balance between active and inactive forms is required for fitness during heat stress(c45). Our results show that, in addition to maintaining fitness during stress, this mode of regulation can be used to control a developmental process.

Nearly all studied members of the HtrA family behave as trimers or multiples thereof(c15–c17, c22, c46, c47). In other systems with two PDZ domains, the first PDZ seems to regulate protease activity directly while the second is thought to mediate rearrangements of core trimers into higher order oligomers(c20, c22, c23). Peptide binding to either the first or the second PDZ domain of MamE can activate proteolysis, although the activation through PDZ2 is much weaker. Furthermore, the protein behaves as a monomer as indicated by gel filtration. Transitions between a monomer and higher order assemblies are rare in the HtrA family, but the positive cooperativity observed for MamE suggests that the active form is indeed a larger assembly(c48). The protein production method described here should enable future structural studies aimed at understanding the potential for novel assembly behavior as well as an unusual regulatory role for the second PDZ domain.

Taken together, our results support the checkpoint model for MamE-dependent proteolysis in regulating the maturation of magnetite crystals. However, the detailed mechanism by which that activity promotes crystal growth remains unclear. One possible mechanism could be by controlling the size of the surrounding membrane. A recent study demonstrated a link between the growth of magnetosome membrane compartments and growth of the magnetite crystals within. This finding led to the proposal that there is a checkpoint regulating a second stage of membrane growth after the onset of biomineralization(c49). In addition to biomineralization defects, proteins normally targeted to the magnetosome membrane are

### Experimental procedures

#### Strains, plasmids and growth conditions

The strains and plasmids used in this study are described in Table 1 and Table 2, respectively. AMB-1 was maintained in MG medium supplemented with kanamycin when necessary as previously described (c6). Magnetic response was measured using the coefficient of magnetism as previously described (c6). Standard molecular biology techniques were used for plasmid manipulation. *E. coli* strains were grown in LB medium supplemented with appropriate antibiotics. Plasmids were maintained in *E. coli* strain DH5α *λpir. E. coli* strain WM3064 was used for plasmid conjugations as described previously(c6).

**Table 1.** Strains used in this study.

| **Strain** | **Organism** | **Description** | **Source** |
| --- | --- | --- | --- |
| AK30 | *M. magneticum* AMB-1 | Wild-type AMB-1 | (6) |
| AK69 | *M. magneticum* AMB-1 | $\Delta mamP$ | (6) |
| AK96 | *M. magneticum* AMB-1 | $\Delta mamE \Delta limE$ | (13) |
| AK94 | *M. magneticum* AMB-1 | $\Delta mamO \Delta R9$ | (13) |
| AK205 | *M. magneticum* AMB-1 | $\Delta mamE \Delta mamO \Delta R9$ | (24) |
| C43 | *E. coli* | protein expression strain | Lucigen |
| BL21<br>CodonPlus | *E. coli* | protein expression strain; Cm<sup>R</sup> | Agilent |
| DH5α ($\lambda pir$) | *E. coli* | standard cloning strain | (6) |
| WM3064 | *E. coli* | mating strain; DAP auxotroph<br>used for plasmid transfer | (6) |

**Table 2.** Plasmids used in this study.

| Plasmid | Description | Source |
| --- | --- | --- |
| pAK605 | Plasmid for expressing protein under control of the *mamH* promoter; non-replicative in AMB-1: integrates upstream of *amb0397* | (24) |
| pAK701 | pAK605 containing N-terminally 3xFLAG tagged *mamE* | This work |
| pAK702 | pAK605 containing C-terminally 3xFLAG tagged *mamE* | This work |
| pAK787 | pAK605 containing N-terminally 3xFLAG tagged *mamP* | This work |
| pAK788 | pAK605 containing C-terminally 3xFLAG tagged *mamP* | This work |
| pAK823 | pAK605 containing N-terminally 3xFLAG tagged *mamO* | (24) |
| pAK1003 | pAK605 containing C-terminally 3xFLAG tagged *mamO* | This work |
| pAK1005 | pLBM4; LIC compatible vector for expressing fusions to the OmpA signal peptide under the control of the *lac* promoter | (28) |
| pAK1004 | pEC86; plasmid constitutively expressing the *E. coli ccm* genes for *c*-type cytochrome maturation | (29) |
| pAK825 | pLBM4 containing 6xHis-TEV-MamE(resi 108-728)-strep fused to the OmpA signal peptide | This work |
| pAK964 | pAK825 with the S297A mutation in MamE | This work |
| pAK619 | pAK605 containing *mamE* | (24) |
| pAK620 | pAK605 containing *mamE*<sup>PD</sup> (H188A T211A S297A mutant) | (24) |
| pAK999 | pAK605 containing *mamE*<sup>QP</sup> (Q294P mutant) | This work |
| pAK1000 | pET based vector expressing the first PDZ domain of MamE (resi 489-588) fused to 6xHis-MBP-TEV at the N-terminus | This work |
| pAK1001 | pET based vector expressing the second PDZ domain of MamE (resi 634-728) fused to 6xHis-MBP-TEV at the N-terminus | This work |
| pAK1002 | pET based vector expressing the first and second PDZ domains of MamE (resi 489-728) fused to 6xHis-MBP-TEV at the N-terminus | This work |

#### Immunoblotting

Whole-cell lysates of AMB-1 strains were prepared from 10mL cultures and analyzed as described previously(c24). For analysis of the autocleavage reaction products, a 1 in 10 dilution of each time-point was separated on a 12% (v/v) acrylamide gel for immunoblotting. The MamE and MamP antibodies have been described previously(24). The anti-6xHis (Sigma), antiFLAG (Sigma), anti-σ70 (Thermo Fisher) and anti-strep (Qiagen) antibodies were purchased from commercial sources.

#### Fractionation of MamO fragments

A strain with the genetic background *AOAR9/FLAG-O* was cultured without shaking at 30 °C in 2 L screw-capped flasks that were filled to the top with MG medium. The cells were harvested by centrifuging at 5k × g for 15 min, redispersed throughout the cytoplasmic membrane in a *mamE* deletion, suggesting an additional defect in magnetosome membrane organization(c6, c13). Magnetosome membrane sizes have not been quantified in this strain. It is tempting to speculate that proteolysis controls a switch that links membrane growth to crystal growth. In this scenario, crystal nucleation in the *E^PD^* cells would not lead to membrane growth, while the *E^QP^* cells would initiate membrane growth before crystals had grown sufficiently, leading to stunted particles in both cases.

suspending in cold 20mL 25mM Tris-HCl pH7.4, re-centrifuging at 8k × g for 10 min and freezing the resulting pellet at −80°C until use. Cell pellets were thawed on ice and re-suspended in 5mL lysis buffer A (10mM Tris-HCl pH 8.0, 50mM NaCl, 1mM EDTA). Pepstatin A and leupeptin were added to a final concentration of 2μg/mL and PMSF was added to 2mM. Lysozyme was added from 50mg/mL stock to a final concentration of 0.5mg/mL and the cells were incubated at room temperature for 15 min. 15mL of lysis buffer B (20mM HEPES-KOH pH7.5, 50mM NaCl, 1. 25mM CaCl_2_) was added along with DTT to 2mM and DNAse to 5μg/mL and the suspension was incubated for 15min at 4°C with agitation. The cells were sonicated twice for 10 seconds and the suspension was centrifuged at 8k × g for 10min to isolate the magnetite-associated material.

The resulting pellet was re-suspended in 5.5 mL solubilization buffer (20mM BisTris-HCl pH7.0, 75mM NaCl, 10% (v/v) glycerol) and CHAPS was added to 1% (w/v) from a 10% (w/v) stock solution. The suspension was incubated at room temperature for 15min with agitation followed by an incubation for 15min at 4°C with agitation. The suspension was centrifuged at 16k × g for 15min. The resulting pellet was re-suspended in 5.5 mL solubilization buffer and the detergent extraction was repeated with 1%(w/v) FosCholine. The FosCholine-soluble material was loaded on a 1mL HiTrap Q FF column (GE Healthcare) that had been equilibrated with solubilization buffer containing 0.03% (w/v) *n*-dodecyl ²-P-D-maltoside (DDM). The column was washed with 10mL of solubilization buffer with 0. 03% (w/v) DDM and eluted with 4mL of buffer Q1 (20mM BisTris-HCl pH7.0, 275mM NaCl, 10% (v/v) glycerol) with 0.03% (w/v) DDM followed by 4mL of buffer Q2 (20mM BisTris-HCl pH7.0, 400mM NaCl, 10% (v/v) glycerol) with 0.03% (w/v) DDM. The Q1 fraction was added to 50μL anti-FLAG M2 resin (Sigma) and incubated at 4°C with agitation for 3 hrs. The resin was isolated by centrifuging at 4k × g and washed with sequential 1mL washes of buffer Q2, buffer Q1 and solubilization buffer each containing 0.03% (w/v) DDM. Bound proteins were eluted by 3 washes with 50μL of 0.2M glycine pH 2.8, which were pooled with 50μL of 1M Tris-HCl pH8.0.

#### Preparation of trypsin digests for liquid chromatography-mass spectrometry (LC-MS)

The concentrated FLAG elution fraction was separated on a 12% (w/v) acrylamide gel and stained with colloidal Coomassie Blue. A ~3 × 10mm section of the gel corresponding to the processed MamO band was excised from the gel and chopped into small pieces. These were washed with 100mM NH4HCO3 followed by reduction and alkylation of cysteines with DTT and iodoacetamide. The gel pieces were then dehydrated by washing with increasing concentrations of acetonitrile in 100mM NH_4_HCO_3_ and dried under vacuum. A 0.1mg/mL solution of trypsin was used to re-swell the gel pieces, and they were incubated overnight at 37°C. The resulting peptides were extracted from the gel slices with successive washes of 0.1% (v/v) formic acid solutions containing increasing concentrations of acetonitrile. The extracts were pooled in a fresh tube, concentrated under vacuum to remove the organic phase and stored at 4°C until analysis.

#### LC-MS

Trypsin-digested protein samples were analyzed using a Thermo Dionex UltiMate3000 RSLCnano liquid chromatograph that was connected in-line with an LTQ-Orbitrap-XL mass spectrometer equipped with a nanoelectrospray ionization (nanoESI) source (Thermo Fisher Scientific, Waltham, MA). The LC was equipped with a C18 analytical column (Acclaim_®_ PepMap RSLC, 150 mm length × 0.075 mm inner diameter, 2 μm particles, 100 Å pores, Thermo) and a 1-μL sample loop. Acetonitrile (Fisher Optima grade, 99.9% (v/v)), formic acid (1-mL ampules, 99+% (v/v), Thermo Pierce), and water purified to a resistivity of 18.2 MΩ-cm (at 25 °C) using a Milli-Q Gradient ultrapure water purification system (Millipore, Billerica, MA) were used to prepare mobile phase solvents. Solvent A was 99.9% water/0.1% (v/v) formic acid and solvent B was 99.9% acetonitrile/0.1% (v/v) formic acid. The elution program consisted of isocratic flow at 2% B for 4 min, a linear gradient to 30% B over 38 min, isocratic flow at 95% B for 6 min, and isocratic flow at 2% B for 12 min, at a flow rate of 300 nL/min.

Full-scan mass spectra were acquired in the positive ion mode over the range *m/z* = 350 to 1800 using the Orbitrap mass analyzer, in profile format, with a mass resolution setting of 60,000 (at *m/z* = 400, measured at full width at half-maximum peak height, FWHM), which provided isotopic resolution for singly and multiply charged peptide ions. Thus, an ion’s mass and charge could be determined independently, i.e., the charge state was determined from the reciprocal of the spacing between adjacent isotope peaks in the *m/z* spectrum. In the data-dependent mode, the eight most intense ions exceeding an intensity threshold of 50,000 counts were selected from each full-scan mass spectrum for tandem mass spectrometry (MS/MS) analysis using collision-induced dissociation (CID). MS/MS spectra were acquired using the linear ion trap, in centroid format, with the following parameters: isolation width 3 *m/z* units, normalized collision energy 30%, default charge state 3+, activation Q 0.25, and activation time 30 ms. Real-time charge state screening was enabled to exclude unassigned and 1+ charge states from MS/MS analysis. Real-time dynamic exclusion was enabled to preclude re-selection of previously analyzed precursor ions, with the following parameters: repeat count 2, repeat duration 10 s, exclusion list size 500, exclusion duration 90 s, and exclusion mass width 20 ppm. Data acquisition was controlled using Xcalibur software (version 2.0.7, Thermo). Raw data were searched against the *Magnetospirillum magneticum* AMB-1 FASTA protein database using Proteome Discoverer software (version 1.3, SEQUEST algorithm, Thermo). Peptide identifications were validated by manual inspection of MS/MS spectra, i.e., to check for the presence of y-type and b-type fragment ions^1^ that identify the peptide sequences(c27).

#### Expression and purification of MamE

In the expression construct used to purify MamE, the N-terminal membrane anchor is replaced with the OmpA signal peptide to produce a soluble protein that can still undergo heme loading in the periplasm(c28). Initial fractionations with an N-terminally 6xHis-tagged form of the protein had significant contamination due to what appeared to be truncated fragments caused by auto-cleavage during expression. To eliminate this problem, a strep tag was added on the C-terminus to allow for a sequential affinity isolation of full-length protein. Finally, MamE has a predicted region of 60-70 disordered residues downstream of the N-terminal membrane anchor and upstream of the trypsin-like domain. Removing this region dramatically improved the solubility

The plasmids pAK825 or pAK964 were transferred to C43 cells (Lucigen) that had been previously transformed with the pEC86 heme-loading plasmid(c29). The transformed cells were maintained at 30°C due to growth defects caused by the plasmids at 37°C. An overnight liquid culture was inoculated into 600mL 2xYT medium supplemented with the appropriate antibiotics. The cultures were grown at 30°C until the OD600 reached ~0.5, at which point the culture was transferred to 20°C. After a 30 min equilibration, the culture was induced with 35μM IPTG. Expression was performed for 12.5-13 hrs at 20°C with shaking at 200 rpm.

Cells were harvested by immediately chilling the cultures on ice and centrifuging at 6k μ g for 10 min. The resulting pellet was resuspended in 50mL of cold osmotic shock buffer (50mM NaPhosphate pH8.0, 1mM EDTA, 20% (w/v) sucrose). Leupeptin (1.5μg/mL), pepstatin A (1.5μg/mL) and lysozyme (0.5mg/mL) were added, and the suspension was rocked at room temperature for 15 min. An equal volume of ice-cold H2O was added and the suspension was rocked on ice for 15 min before centrifuging at 8k × g for 10 min to remove debris.

The resulting supernatant was added to 3 mL Ni-NTA resin (Qiagen) and supplemented with NaCl (150mM), DNAseI (5μg/mL), NP-40 (0.1% (v/v)) and MgCl_2_ (2.5mM). The slurry was rocked at 4°C for 30 min and the beads were allowed to settle. After decanting the upper phase, the slurry was poured into a column, washed with 10 column volumes of Ni wash buffer (25mM Tris-HCl pH7.4, 250mM NaCl, 10mM imidazole, 10% (v/v) glycerol) and the bound proteins eluted with Ni elution buffer (25mM Tris-HCl pH7.4, 250mM NaCl, 250mM imidazole, 10% (v/v) glycerol). The Ni-NTA eluent was loaded onto a 1mL StrepTrap column (GE Healthcare), which was then washed with 5mL strep wash buffer (25mM Tris-HCl pH7.4, 250mM NaCl, 10% (v/v) glycerol). Bound proteins were eluted in strep wash buffer containing 2.5mM desthiobiotin. The purified protein was concentrated in 50kDa cutoff ultrafilter while simultaneously removing the desthiobiotin by repeated dilution and concentration with strep wash buffer. The concentration was determined by the Bradford method using bovine serum albumin to prepare a standard curve. Aliquots were frozen in liquid N_2_ and stored at −80°C until use.

#### Analysis of MamO1 peptide cleavage

A custom peptide with the sequence 5carboxymethylfluorescein-Thr-Gln-Thr-Val-Ala-Ala-Gly-Ser-Lys(CPQ2)-D-Arg-D-Arg was obtained commercially (CPC Scientific). The peptide was dissolved in DMSO and stored at −20°C. 5X substrate solutions with various concentrations of the MamO1 peptide were prepared in assay buffer (50mM Tris-HCl pH8.0) containing 0.05% (v/v) NP-40 and 1.6% (v/v) DMSO. To initiate the reaction, 10μL samples of the substrate mix were added to 40μL of MamE protein solution that had been diluted to 125nM in assay buffer in a 96-well plate. The fluorescence was scanned (excitation: 485nm; emission 538nm) every 5 min for 2 hrs in a Tecan plate reader.

The slope was determined from the linear portion of each reaction. Cleavage rates were calculated by making a standard curve from a MamO 1 cleavage reaction that had been incubated for 24hrs to allow for complete hydrolysis. Specific activities were determined by normalizing these cleavage rates to the enzyme concentration. Rates were plotted as a function of peptide concentration and fit to the Hill form of the Michaelis-Menten equation using the Kaleidagraph software package:

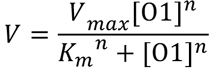

#### Analysis of MamE auto-proteolysis

25μL reactions were prepared by adding 1μL of activating peptide dissolved in DMSO at the appropriate concentration to 24μL of MamE diluted to 2μM in assay buffer. The reactions were incubated at 30°C, and 8μL aliquots were removed at the appropriate times. Each aliquot was quenched immediately by mixing with SDS sample buffer. Equal volumes of each aliquot were separated on a 12% (w/v) acrylamide SDS-PAGE gel and stained with Coomassie Blue to visualize the processing pattern.

#### Expression and purification of PDZ Domains

For all three PDZ domain constructs (EP1, EP2 and EP12), the appropriate plasmids for expression as N-terminal 6xHis-MBP-TEV fusions were transformed into BL21 Codon Plus cells. Cultures were grown in 2xYT at 37°C until the OD_600_ reached ~0.8 at which point they were transferred to 20°C for 30min followed by induction with 0.1mM IPTG and expression overnight. The cells were harvested by centrifugation, resuspended in resuspension buffer (25mM Tris-HCl pH7.4, 800mM NaCl, 10mM imidazole, 10% (v/v) glycerol) and frozen at −80°C until use.

For protein purification, the cells were thawed on ice and sonicated for three 30-second cycles. The lysate was clarified by centrifuging at 13k × g for 30 min. The resulting supernatant was loaded on a 3mL Ni-NTA column that had been equilibrated in resuspension buffer. After washing with 10 column volumes of resuspension buffer and 10 column volumes of wash buffer 2 (25mM Tris-HCl pH7.4, 400mM NaCl, 25mM imidazole, 10% (v/v) glycerol), bound proteins were eluted with Ni elution buffer.

For the purification of the EP1 and EP2 proteins, the elution fractions were dialyzed overnight against AEX buffer A (25mM BisTris-HCl pH7.0, 75mM NaCl and 10% (v/v) glycerol). The desalted protein was passed through a 1mL HiTrap QFF column (GE Healthcare) and the flow-through was concentrated in a 50kDa cutoff ultrafilter, injected onto a16/60 Superdex 200 column and developed in Storage Buffer. Each protein eluted as a single symmetrical peak. The peak fractions were concentrated in a 50kDa ultrafilter and small aliquots were frozen in liquid N2 and stored at −80°C for use in the phage display experiments.

For the purification of the EP12 protein, the elution fraction was dialyzed overnight against digest buffer (50mM NaPhosphate pH8.0, 75mM NaCl, 5mM imidazole, 10% (v/v) glycerol) in the presence of 6xHis tagged TEV protease to remove the 6xHis-MBP tag. The resulting sample was passed through a 3mL Ni-NTA column that had been equilibrated in digest buffer. The flowthrough fraction was concentrated in a 10kDa ultrafilter, passed through a 1mL HiTrap SP FF column and concentrated again before injection on a16/60 Superdex 200 column that was developed in storage buffer. The protein eluted as a single symmetrical peak. The peak fractions were concentrated in a 10kDa ultrafilter and small aliquots were frozen in liquid N2 and stored at −80°C for use in the fluorescence anisotropy experiments.

#### Phage display

C-terminally and N-terminally displayed peptide libraries were used to assess the peptide binding preferences of MamE PDZ1 and PDZ2. The C-terminal peptide library consisted of random decapeptides constructed using 10 consecutive NNK degenerate codons encoding for all 20 natural amino acids and fused to the C terminus of a mutant M13 bacteriophage major coat protein (2 × 10^10^ unique members)(c30, c31). The N-terminal peptide library consisted of random hexadecapeptides constructed using 16 consecutive mixes of 19 codon trimers (cysteine and STOP codons were excluded) and fused to the N terminus phage coat protein (2.4 × 10^11^ unique members)(c32).

The phage display selections followed an established protocol used previously for the PDZ human domains(c33, c34). Briefly, each library was separately cycled through rounds of binding selection against each immobilized MBP-PDZ fusion protein on 96-well MaxiSorp immunoplates (Nalge Nunc). After each round, phage were propagated in *E. coli* XL1-blue cells (Stratagene) supplemented with M13-KO7 helper phage (New England Biolabs) to facilitate phage production. After four rounds of selection, phages from individual clones were analyzed in a phage enzyme-linked immunosorbent assay (ELISA). Phages that bound to the MBP-PDZ fusion but not a control MBP were subjected to DNA sequence analysis.

#### Fluorescence anisotropy

Peptides with the following sequences were synthesized commercially with a fluorescein-aminocaproic acid group fused to the N-terminus: WSQEMEDWFWQMPLSG (PDZ1^*^) and MEDYGIFMTSPEGPWA (PDZ2^*^). Each peptide was diluted to a concentration of 40nM in 25mM Tris-HCl pH7.4 containing 0.25mg/mL bovine serine albumin. A dilution series of EP12 protein was prepared in storage buffer. 6μL of the ligand solution was added to 18μL of the appropriate protein solution in a 384-well plate, and the mixture was allowed to equilibrate at room temperature for 15 min. Polarization measurements were made at 535nm using a Perkin Elmer Victor 3 V 1420 plate reader. The resulting anisotropy values were plotted as a function of protein concentration and fit to a single site binding model using the Kaleidagraph software package:

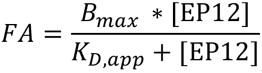

## Acknowledgements

This work was supported by grants from the National Institutes of Health (R01GM084122) and the National Science Foundation (1504610) to AK. The QB3/Chemistry Mass Spectrometry Facility at the University of California, Berkeley receives support from the National Institutes of Health (1S10OD020062-01).

## Conflict of Interest

The authors declare no conflict of interest.

